# Loss of FAS1 function reveals rescue of an aberrant intra-S-phase checkpoint by the G2/M checkpoint regulator SOG1^1[OPEN]^

**DOI:** 10.1101/2021.02.22.432330

**Authors:** Thomas Eekhout, Martina Dvorackova, José Antonio Pedroza Garcia, Martina Nespor Dadejova, Pooneh Kalhorzadeh, Hilde Van den Daele, Ilse Vercauteren, Jiri Fajkus, Lieven De Veylder

## Abstract

The WEE1 and ATR kinases represent important regulators of the plant intra-S-phase checkpoint, as evidenced by the hypersensitivity of *WEE1*^KO^ and *ATR^KO^* roots to replication inhibitory drugs. Here, we report on the identification of a defective allele of the *FASCIATA1* (*FAS1*) subunit of the chromatin assembly factor 1 (CAF-1) complex as a suppressor of *WEE1*- or ATR-deficient plants. We demonstrate that lack of *FAS1* activity results in the activation of an ATM- and SOG1-mediated G2/M-arrest that makes the ATR and WEE1 checkpoint regulators redundant. This ATM activation accounts for telomere erosion and loss of ribosomal DNA described for the *fas1* plants. Knocking out *SOG1* in the *fas1 wee1* background restores replication stress sensitivity, demonstrating that SOG1 plays a prominent role as secondary checkpoint regulator in plants that fail to activate the intra-S-phase checkpoint.

**One-Sentence Summary:** Lack of the chromatin assembly factor-1 subunit FAS1 results in a DNA damage response that overrules the need for replication checkpoint activators.

## INTRODUCTION

Plants can be exposed to adverse conditions such as drought, heat, and UV radiation. Many of these stresses result in the generation of reactive oxygen species (ROS), which can oxidize and damage the DNA (Roldán-Arjona and Ariza, 2009). The response to this DNA damage is dependent on the type of damage, with double-strand breaks (DSBs) predominantly activating the ATAXIA TELANGIECTASIA MUTATED (ATM) kinase, whereas the accumulation of single-stranded DNA (ssDNA), arising due to problems during the DNA replication process, activates the ATM AND RAD3-RELATED (ATR) kinase. In mammals, the checkpoint kinases Chk1 and Chk2 and the transcription factor p53 operate downstream of ATM/ATR, governing transcriptional responses to DNA damage (Zhou and Elledge, 2000). In plants however, Chk1 and Chk2 kinases are absent, as is a homolog of p53. Rather, the functionally similar but structurally unrelated transcription factor SUPPRESSOR OF GAMMA-RESPONSE 1 (SOG1) governs most transcriptional responses to DNA damage in plants (Yoshiyama et al., 2009). When plants sense DNA damage, the ATM and ATR kinases phosphorylate SOG1 and in this way activate its transcriptional activity (Yoshiyama et al., 2013; Sjogren et al., 2015), which then induces expression of both DNA repair and cell cycle inhibitory genes (Bourbousse et al., 2018; Ogita et al., 2018).

During the S-phase of the cell cycle, the DNA is unpacked from its protective chromatin wrapping and uncoiled to allow passage of the replication fork complex. In the event of problems during the replication process, DNA integrity checkpoints are activated that slow down the cell cycle to grant the cell time to repair the damage. A major player in the plant S-phase checkpoint is the WEE1 kinase, which phosphorylates and, in this way, inhibits activity of the drivers of the cell cycle, being the cyclin-dependent kinase (CDK)/cyclin complexes (De Schutter et al., 2007). For Arabidopsis (*Arabidopsis thaliana*), it has been shown that WEE1 is activated downstream of the ATR kinase when replication stress occurs (De Schutter et al., 2007). The kinase plays an essential role to survive replication stress, since *WEE1*^KO^ plants show a hypersensitive response to the replication inhibitory drug hydroxyurea (HU) that lowers the available dNTP pool (De Schutter et al., 2007; Cools et al., 2011). Through an EMS mutagenesis suppressor screen, previous work has identified two subunits of the RNase H2 complex that, when mutated, rescue the hypersensitive response of the *WEE1*^KO^ plants to HU, likely through allowing to substitute dNTP by rNTPs into the replicating DNA (Kalhorzadeh et al., 2014; Eekhout et al., 2015), again suggesting that WEE1 activity is activated in response to a depletion of the available dNTP pool. More recently rescue of the *WEE1*^KO^ plants grown on HU has been reported for mutants in the RNA splicing factor factors PRL1 and CDC5, and the F-box like 17 (FBL17) E3-ubiquitine ligase, where the HU induced cell cycle arrest activated by WEE1 has been attributed to incorrect splicing of cyclin mRNA transcripts and stabilization of CDK inhibitory proteins, respectively (Pan et al., 2021; Wang et al., 2021).

When DNA is replicated, new histones need to be added to the nascent DNA and epigenetic marks need to be copied from the parental chromatin to the newly assembled one. Histones H3 and H4 are bound as a dimer by the histone chaperone ASF1 and in this way imported into the nucleus (Zhu et al., 2011). There they are transferred to the chromatin assembly factor-1 (CAF-1) complex, which consists of three subunits, called CAC1-3 in budding yeast (Kaufman et al., 1997), p155, p60 and p48 in human (Smith and Stillman, 1989; Kaufman et al., 1995) and FASCIATA1 (FAS1), FASCIATA2 (FAS2) and MULTICOPY SUPPRESSOR OF IRA 1 (MSI1) in plants (Kaya et al., 2001). The CAF-1 complex is responsible for both the formation of (H3-H4)_2_ tetramers from newly synthesized H3-H4 histone dimers, and the association of these tetramers with synthesized DNA through CAF-1 interaction with the PROLIFERATING CELL NUCLEAR ANTIGEN (PCNA) complex (Smith and Stillman, 1989; Kaufman et al., 1995; Verreault et al., 1996). After loading of the histone (H3-H4)_2_ tetramers onto the DNA, histones H2A and H2B can be added by NUCLEOSOME ASSEMBLY PROTEIN1 (NAP1) and NAP1-RELATED PROTEIN NRP chaperones (Zhu et al., 2006).

FASCIATA mutants were first described as plants with thickened and flattened stems, and abnormal phyllotaxy (Leyser and Furner, 1992). Further research showed that the CAF-1 complex is needed to maintain cellular organization in the meristems, since *fas1* plants display disorganized meristems and aberrant expression of stem cell regulators such as *WUSCHEL* and *SCARECROW* (Kaya et al., 2001). Absence of CAF-1 function has additionally been linked to various changes in cell cycle progression, including an extended S-phase (Schönrock et al., 2006) and decreased CDKA;1 activity (Ramirez-Parra and Gutierrez, 2007). Consistently, *fas1* mutants show an altered distribution of cells with increased ploidy levels due to enhanced endoreplication (Endo et al., 2006; Exner et al., 2006; Jacob et al., 2014), which is an adaptation of the cell cycle when the cell skips mitosis but increases the DNA content. Knocking out *ATM* partially suppresses the decrease in cell number and the increase in ploidy levels seen in the *fas1* mutant (Hisanaga et al., 2013). CAF-1 also suppresses homologous recombination (Endo et al., 2006; Kirik et al., 2006), and its deletion leads to the constitutive activation of DNA repair genes (Schönrock et al., 2006; Hisanaga et al., 2013). *fas1 atm* double mutants lack this constitutive induction of *RAD51*, *BRCA1* and two *PARP* genes, being typical DSB-responsive targets, indicating that ATM plays an essential role in relaying the DNA damage response in *fas1* (Hisanaga et al., 2013). In addition to growth phenotypes, *fas1* mutants show a progressive shortening of telomeres and loss of 45S rDNA as plants go through mitosis (Mozgová et al., 2010), correlated with RAD51B-mediated homology-dependent repair (Muchová et al., 2015).

In this work, we describe that a mutation in the *FAS1* gene rescues the hypersensitivity of *WEE1*^KO^ plants to HU. We demonstrate that the *fas1* mutation eliminates the need for WEE1 as S-phase checkpoint regulator, likely because of activation of a G2/M checkpoint controlled by ATM and SOG1. Accordingly, we show that the loss of rDNA and telomere shortening in *fas1* results from ATM activation. Finally, we show that the checkpoint responsible for the rescue of HU sensitivity in *fas1 wee1* plants is dependent on SOG1.

## RESULTS

### A Mutation in the FAS1 Subunit of the CAF-1 Complex Rescues HU Sensitivity of *WEE1*^KO^ Plants

*WEE1*^KO^ seedlings (*wee1-1*) show a root growth arrest when grown on HU-containing medium (De Schutter et al., 2007). This phenotype was used to identify suppressor mutations within an ethyl methanesulfonate (EMS) mutagenized M2 *wee1-1* population, screening for restoration of root growth on 0.75 mM HU. One of the identified mutants was line 49-2, which showed a partial recovery of root growth on HU. To map the mutation responsible for this phenotype, we used an advanced backcrossing strategy to introduce the *wee1-1* allele into a Landsberg *erecta* background (see Materials and Methods). After four generations of backcrossing, the *wee1-1* (L*er*) line contained at least 98% of L*er* DNA as tested by AFLP. This *wee1-1* (L*er*) line was subsequently crossed to line 49-2, after which the segregating F2 population was used for bulked segregant next-generation sequencing-based gene mapping using the SHOREmap algorithm (Schneeberger et al., 2009). The underlying mutation in line 49-2 was pinpointed to a base pair change in codon 281 of the *At1g65470* gene, resulting in the generation of a premature stop codon. This base pair change was confirmed in the original 49-2 line by direct Sanger sequencing (Supplemental Fig. S1). The *At1g65470* gene is annotated as the p150 subunit of the chromatin assembly factor-1 (CAF-1) complex, also known as *FAS1*. To confirm the loss of FAS1 activity, we crossed the *wee1-1* mutation out of line 49-2 and compared the resulting single *fas1* mutant with an available T-DNA insertion knockout line (GABI-095A01), referred to as *fas1-8*, by analyzing described phenotypes (Supplemental Fig. S2). Ploidy levels were increased in line 49-2 to similar levels as observed for *fas1-8*, while leaf size of 21-d-old plants was equally decreased in both lines compared to Col-0 (Supplemental Fig. S2, B and C). Moreover, the *fas1-8* line has a similar rosette phenotype as line 49-2 and other described *fas1* alleles, with the leaves being smaller and serrated (Supplemental Fig. S2D). Subsequently, the *fas1-8* allele was crossed into the *wee1-1* background and tested for HU sensitivity (Fig. 1A-C). Because root growth analysis demonstrated a similar rescue of the *wee1-1* phenotype as the 49-2 line when grown on HU-containing medium, the *fas1-8* line was predominantly used for further experiments to avoid potential effects of secondary mutations in line 49-2 caused by the EMS treatment.

**Figure 1.**
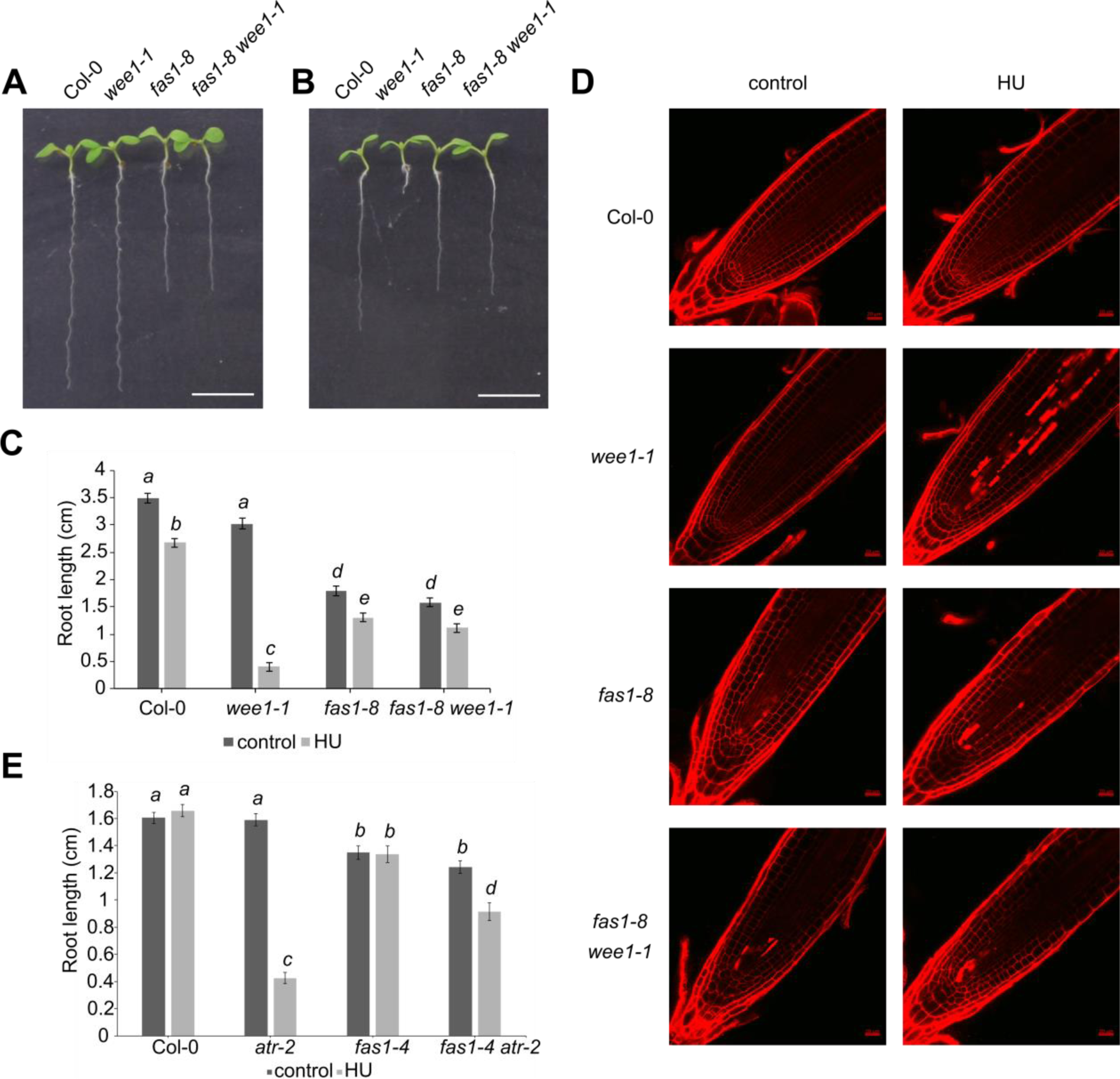
A mutation in *FAS1* partially rescues the *wee1-1* hypersensitive phenotype on hydroxyurea. (A) and (B) Root growth of 7-d-old WT (Col-0) and *wee1-1, fas1-8* and *fas1-8 wee1-1* plants grown on control medium (A) or medium supplemented with 0.75 mM HU (B). Bar = 1 cm. (C) Quantification of the root growth shown in (A) and (B). Data represent mean ± SEM (n > 10). Significance was tested with mixed model analysis. Means with different letters are significantly different (p < 0.05). (D) Confocal microscopy images of root meristem of 7-d-old WT (Col-0) and *wee1-1, fas1-8* and *fas1-8 wee1-1* plants transferred for 24 h to control medium or medium supplemented with 1 mM HU and stained with propidium iodide. Bar = 20 µm. (E) Quantification of root growth of Col-0, *atr-2, fas1-4* and *fas1-4 atr-2* at 7 DAS grown on medium containing -/+ 0.25 mM HU. Data represent mean ± SEM (n > 10). Significance was tested with mixed model analysis. Means with different letters are significantly different (p < 0.05).

### WEE1 Function Is Redundant in *fas1-8* Plants Due to Secondary Checkpoint Activation

Strikingly, root growth did not differ significantly between *fas1-8* and *fas1-8 wee1-1* plants when grown on HU, indicating that introgression of *wee1-1* into the *fas1-8* background did not enhance HU sensitivity of *fas1-8* (Fig. 1B). Additionally, microscopic analysis of the root meristem revealed that *fas1-8 wee1-1* reverted the severe cell death phenotype that is observed in roots of *wee1-1* plants when grown in the presence of HU, but more importantly did not show more cell death compared to the *fas1-8* single mutants (Fig. 1D). We therefore hypothesized that a secondary cell cycle checkpoint might be activated in the *fas1-8 wee1-1* plants, making the WEE1-regulated S-phase checkpoint redundant.

If the checkpoint activated in *fas1* plants can overrule the sensitivity of *WEE1*^KO^ plants to HU, it is expected to also rescue the absence of *ATR*, which is operating upstream of *WEE1*, as demonstrated by a similar HU hypersensitivity phenotype of *atr* mutant plants (Culligan et al., 2004). Indeed, when growing *fas1-4 atr* plants on HU-containing medium, a partial rescue of root growth was observed compared to the *atr* plants (Fig. 1E), confirming our hypothesis that the checkpoint activated by the absence of *FAS1* acts dominantly over the ATR-mediated intra-S-phase checkpoint.

To investigate the nature of the checkpoint being activated by absence of *FAS1*, we performed 5-ethynyl-2’-deoxyuridine (EdU) staining to measure S-phase and total cell cycle duration (Hayashi et al., 2013). Despite the anticipated role of FAS1 during the replication process, S-phase duration was not or only slightly extended in *fas1-8* and *fas1-8 wee1-1*, respectively, under control growth conditions (Fig. 2A and Supplemental Table S1). In contrast, a dramatic increase (> 10 h) in total cell cycle length was observed (Fig. 2B and Supplemental Table S1). An EdU pulse chase experiment displayed a > 4-h delay of replicating cells to enter the M phase (Fig. 2C), indicating that the root growth phenotype of *fas1-8* and *fas1-8 wee1-1* plants is likely due to an extended G2 phase, corroborating a similar conclusion drawn before based on an RT-PCR analysis using cell cycle reporter genes (Hisanaga et al., 2013). To understand the rescue mechanism of the *wee1* phenotype on HU, we similarly performed an EdU time series on plants transferred to medium containing HU. As expected, S-phase length in Col-0 was increased in a WEE1-dependent manner when grown on HU (Figure 2A and Supplemental Table S2). Additionally, HU-induced replication stress increased both S-phase and the total cell cycle length in the *fas1-8* mutant. The *fas1-8 wee1-1* double mutant, in comparison, appeared insensitive to the HU-induced replication stress, because the duration of both the S-phase and the total cell cycle was not further changed compared to the untreated plants (Fig. 2, A and B). It suggests that similarly to what is observed in wild-type (WT) plants, in *fas1-8* the extended S-phase during replication stress occurs in a WEE1-dependent manner. However, in both *fas1-8* and *fas1-8 wee1-1*, the increase in total cell cycle length (being > 9 h compared to Col-0 plants) far exceeds that of the increase in S-phase duration (being maximum > 2.4 h compared to Col-0 plants), suggesting that also in the presence of HU, the root growth phenotype of both mutants is predominantly due to a G2/M cell cycle arrest.

**Figure 2.**
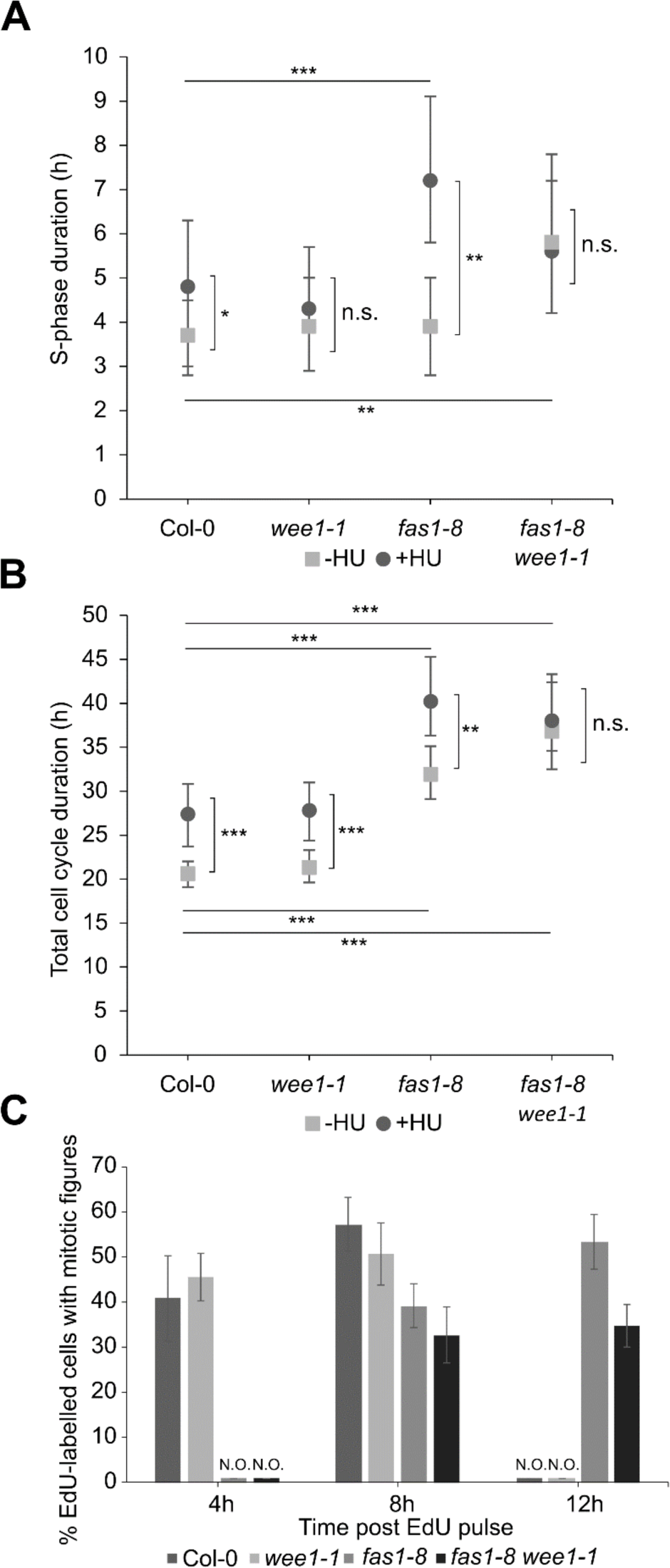
Cell cycle parameters of Col-0, *wee1-1, fas1-8* and *fas1-8 wee1-1* root tip cells under control conditions and treated with HU. (A) and (B) S-phase (A) and total cell cycle (B) duration were measured using a time course of EdU staining according to the protocol of Hayashi et al. (2013). Data represent mean ± 95% confidence intervals (n.s., not significant; *, p < 0.05; **, p < 0.01; ***, p < 0.001). For S-phase duration, comparisons were tested using ANOVA with Tukey correction. Total cell cycle duration was tested using ANOVA with F-tests to statistically test the equality of means (n > 5). (C) EdU pulse-chase experiment. Mitotic figures containing EdU signal were counted in Col-0, *wee1-1*, *fas1-8* and *fas1-8 wee1-1* nuclei at 4 h, 8 h and 12 h after a short EdU pulse. Data represent percentage of EdU-labelled cells among mitotic figures (n > 5).

### ATM Predominantly Accounts for the Observed G2/M Arrest in *fas1*

Multiple transcriptome studies of *fasciata* mutants have been published (Schönrock et al., 2006; Hisanaga et al., 2013; Mozgová et al., 2015), each using different mutant backgrounds and tissues. Meta-analysis of the different available datasets (see methods) allowed us to identify a core set of genes that are, compared to control plants, specifically upregulated as a result of the absence of a functional CAF-1 complex. This meta-analysis yielded 28 genes, being induced in at least two of the three datasets (Fig. 3A). Among these, 15 were previously pinpointed as DNA damage response (DDR) hallmark genes (Fig. 3B) (Yi et al., 2014), corresponding with a statistically significant enrichment (p < 0.001, hypergeometric test), indicating that regardless of the developmental context or affected subunit, dysfunction of the CAF-1 complex leads to activation of the DNA damage response (Fig. 3C).

**Figure 3.**
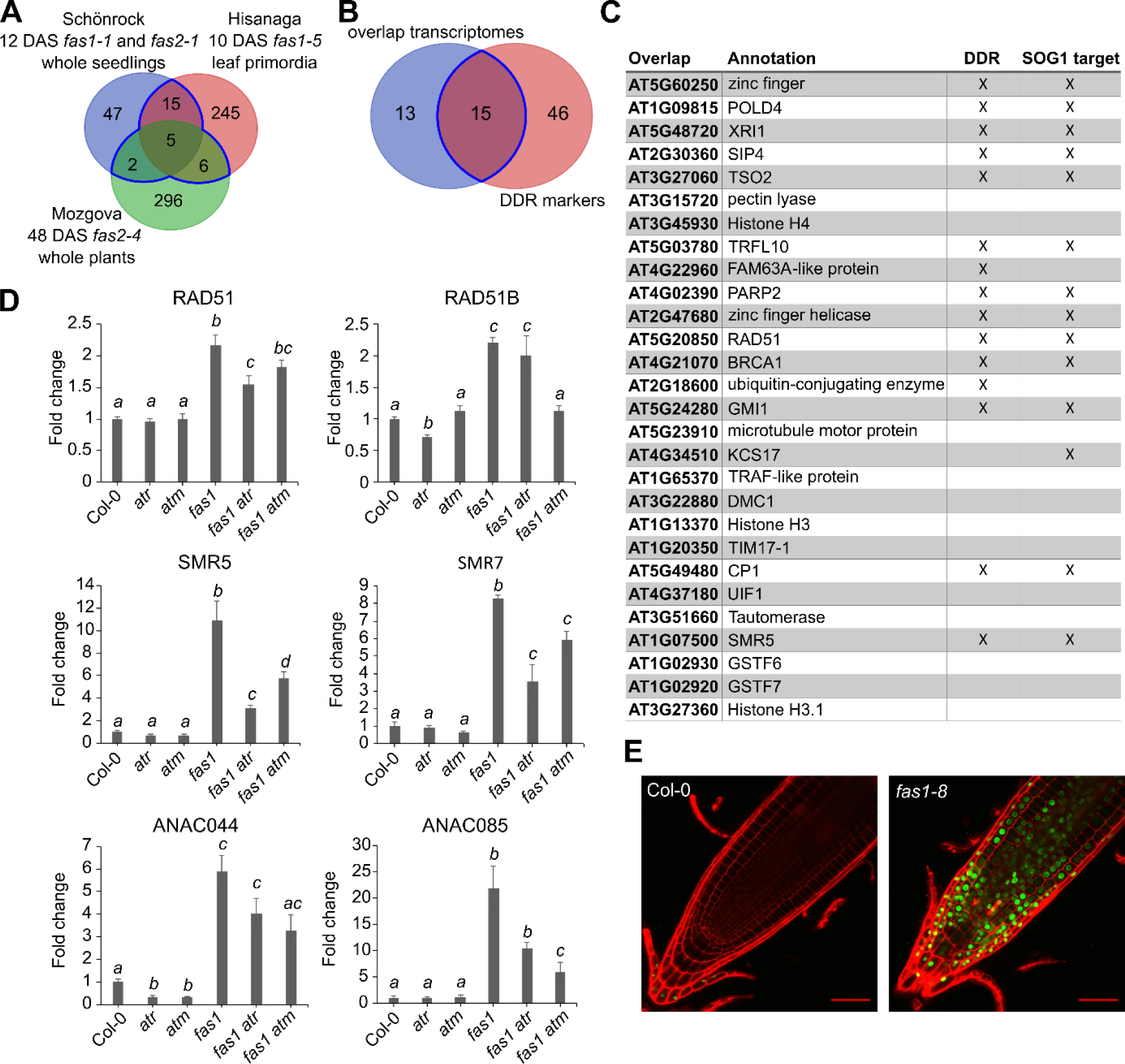
CAF-1 dysfunction leads to the activation of the DDR genes. (A) Overlap of upregulated genes in published transcriptome datasets of *CAF-1* mutants (Schönrock et al., 2006; Hisanaga et al., 2013; Mozgová et al., 2015). (B) Overlap of the core set of upregulated genes in the CAF-1 datasets and the list of DDR markers (Yi et al., 2014). (C) List of upregulated genes in the overlap shown in (A), with SOG1 targets as defined by Bourbousse et al. (2018). (D) Relative expression levels in root tips of 7-d-old WT (Col-0), *atr-2*, *atm-2, fas1-4, fas1-4 atr-2* and *fas1-4 atm-2* mutant plants as determined by qRT-PCR. Data represent mean ± SEM, normalized to WT levels that were arbitrarily set to one (n = 3). Significance was tested through Student’s *t*-test. Means with different letters are significantly different (p < 0.05). (E) Expression of a GFP-reporter construct for SMR7 in Col-0 and *fas1-8* background. Roots were stained with propidium iodide. Bar = 50 µm.

To investigate the dependency of the upregulated DDR genes on the S-phase or G2/M-phase checkpoint, we tested expression of representative genes, both from the meta-analysis and close orthologs of these, involved in DNA repair and checkpoint regulation in both *fas1-4 atr-2* and *fas1-4 atm-*2. Whereas induction of *RAD51* appeared to depend on both ATM and ATR, induction of *RAD51B*, which is involved in single-strand annealing (Serra et al., 2013), was only dependent on ATM (Fig. 3D). Expression of the cell cycle checkpoint regulators *SMR5* and *SMR7* is dependent on both ATM and ATR (Fig. 3D), implying that both DDR kinases play a role in some aspects of the transcriptional response of *fas1* plants. Confocal images of an *SMR7:GUS-GFP* reporter line clearly showed induction of this gene in the *fas1* background throughout the whole root meristem (Fig. 3E). The ANAC044 and ANAC085 transcription factors, demonstrated to account for a G2/M cell cycle arrest in response to DNA damage (Takahashi et al., 2019), were strongly upregulated in the *fas1* mutant and seem to be regulated by both ATM and, to a lesser extent, ATR (Fig. 3D).

Given the role of FAS1 in chromatin assembly, the constitutive higher expression level of DDR genes might be linked to an altered chromatin state rather than inflicted DNA damage. In such case, applying DNA damage to the *FAS1* mutant plants might result in a hyper-induction of DDR gene expression. To test this hypothesis, the different genotypes described were treated with bleomycin, a chemical that causes DSBs, and expression levels of *RAD51* and *SMR7* were measured as examples of DNA repair and cell cycle regulation, respectively. For both, bleomycin treatment upregulated expression in the *fas1* background to a level comparable to that of treated control plants (Supplemental Fig. S3) that remained dependent on both ATM and ATR, confirming that the upregulation of these genes in the *fas1* mutant is primarily caused by activation of the DDR by ATM and ATR.

To understand the influence of ATM and ATR on cell cycle progression in the *fas1* mutant, we calculated S-phase and total cell cycle duration in the *fas1-4* versus *fas1-4 atm-2* and *fas1-4 atr-2* mutants based on EdU time courses. Both *fas1-4 atm-2* and *fas1-4 atr-2* displayed a shorter cell cycle duration compared to *fas1-4* (Fig. 4B and Supplemental Table S3), suggesting that both ATM and ATR contribute to the cell cycle arrest. Only in *fas1-4 atm-2*, the increase in cell cycle duration was accompanied with an increase in S-phase duration (Fig. 4A and Supplemental Table S3), likely due to ATR activity. The lack of an increased S-phase duration in *fas1-4 atr-2*, in contrast, suggests that ATM activity predominantly results in an extended G2 phase.

**Figure 4.**
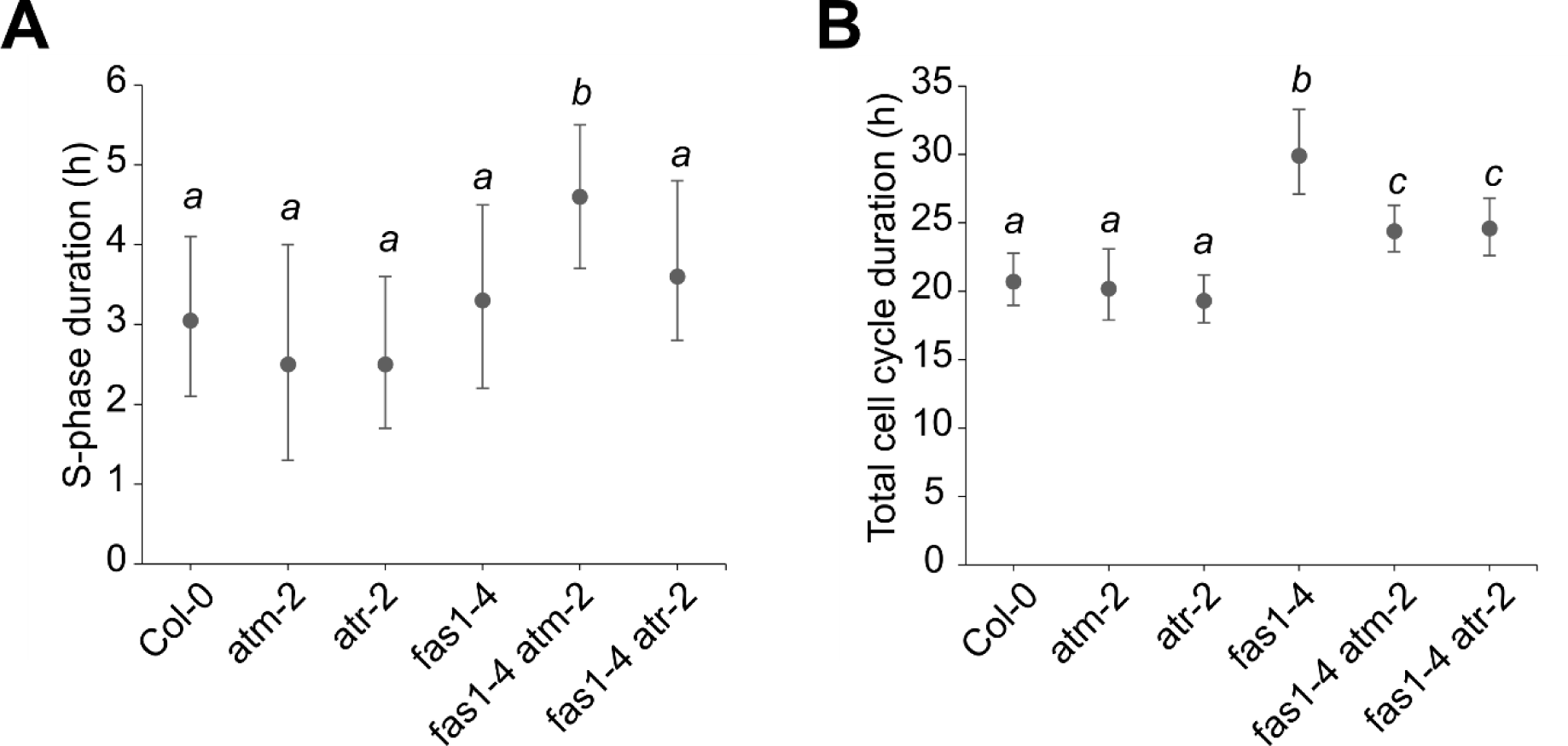
Influence of ATM and ATR on the checkpoint activated in *fas1* plants. (B) and (C) S-phase duration (B) and total cell cycle (C) duration of Col-0, *atm-2*, *atr-2 fas1*- *4*, *fas1-4 atm-2* and *fas1-4 atr-2*, as determined by EdU time course according to the protocol of Hayashi et al. (2013). Data represent mean ± 95% confidence intervals. Means with different letters are significantly different (p < 0.05). For S-phase duration, comparisons were tested using ANOVA with Tukey correction. Total cell cycle duration was tested using ANOVA with F-tests to statistically test the equality of means (n > 5).

If the DSBs caused by the *fas1* mutant background are the cause of the ATM-dependent G2 checkpoint that rescues *wee1-1* plants on replication stress, then chemicals that induce DSBs such as zeocin should equally be able to rescue *wee1-1* plants grown on HU. To test this hypothesis, we grew Col-0 and *wee1-1* plants on either zeocin, HU, or a combination of the two and quantified root growth. As expected, the combination of zeocin and HU was able to partially rescue the *wee1-1* root growth phenotype (Fig 5).

**Figure 5.**
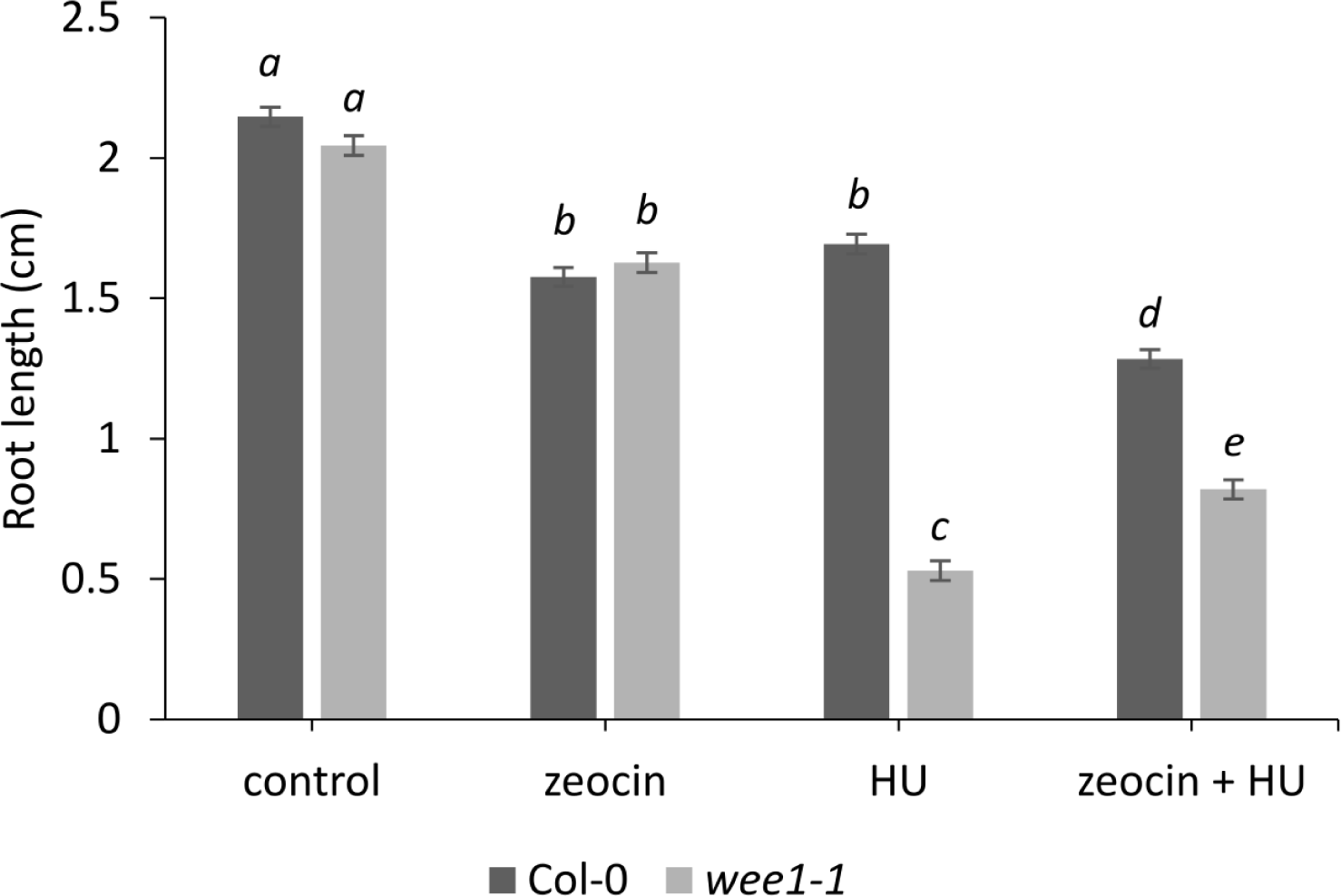
Zeocin partially rescues the *wee1-1* hypersensitive phenotype on hydroxyurea. Quantification of root growth of 7-d-old WT (Col-0) and *wee1-1* plants grown on control medium or medium supplemented with 0.5 mM HU, 12 µM zeocin or both. Data represent mean ± SEM (n = 3, with > 9 plants in each repeat). Significance was tested with mixed model analysis. Means with different letters are significantly different (p < 0.05).

### SOG1 Is Accountable for the Rescue of the *fas1 wee1* Root Growth Phenotype Induced by Replication Stress

To analyze whether the G2/M checkpoint activated by ATM accounts for the HU-resistant phenotype of the *fas1 wee1* mutant plants, *fas1-4 wee1-1 atm-2* triple mutant plants were generated. Root growth was measured on control medium and medium supplemented with HU. To avoid the pleiotropic effects of the *fas1* background (Mozgova et al., 2010; Mozgova et al., 2018), a population homozygous for *atm-2 wee1-1* and segregating for *fas1* was used, after which plants were genotyped for the *fas1* mutation. Under control conditions, *fas1-4 wee1-1 atm-2* roots are indistinguishable from WT plants, corroborating the evidence that activation of ATM causes the root growth penalty in *fas1* mutants. When grown under replication stress, the *fas1-4 wee1-1 atm-2* plants became slightly more sensitive, but were still more resistant than *wee1-1* plants (Fig. 6A). This indicates that in the absence of ATM, ATR might partially contribute to the checkpoint activation.

**Figure 6.**
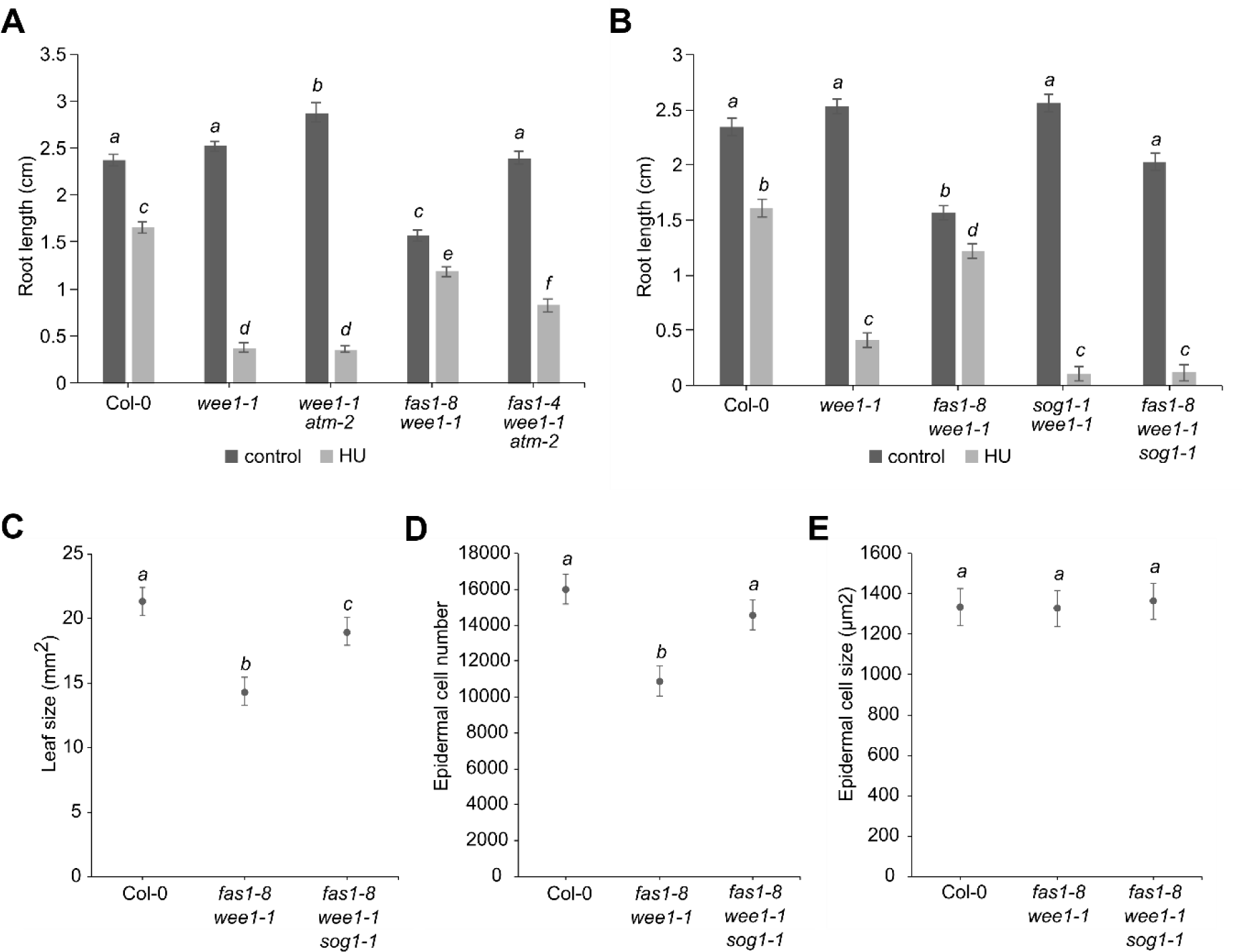
Rescue of *fas1 wee1* on replication stress is dependent on SOG1 but only partially on ATM. (A) Quantification of root growth of Col-0, *wee1-1, wee1-1 atm-2*, *fas1-8 wee1-1* and *fas1-4 wee1-1 atm-2* at 7 DAS grown on medium containing -/+ 0.75 mM HU. Data represent mean ± SEM (n > 10). Significance was tested with mixed model analysis. Means with different letters are significantly different (p < 0.05). (B) Quantification of root growth of Col-0, *wee1-1, fas1-8 wee1-1, sog1-1 wee1-1* and *fas1-8 sog1-1 wee1-1* at 7 DAS grown on medium containing -/+ 0.75mM HU. Data represent mean ± SEM (n > 10). Significance was tested with mixed model analysis. Means with different letters are significantly different (p < 0.05). (C-E) Leaf size (C), epidermal cell number (D) and cell size (E) of first leaves at 21 DAS of Col-0, *fas1-8 wee1-1*, and *fas1-8 sog1-1 wee1-1*. Data represent mean ± SEM (n = 5, with 2 leaves per repeat). Means with different letters are significantly different (p < 0.05).

Downstream of both ATM and ATR, the SOG1 transcription factor can be found. Correspondingly, 14 out of the 28 genes identified in the meta-analysis of *fasciata* transcriptomes were found to be direct SOG1 target genes (Fig. 3C) (Bourbousse et al., 2018; Ogita et al., 2018). Therefore, we generated *fas1-8 wee1-1 sog1-1* plants and compared them with *fas1-8 wee1-1* plants of the same generation. Under control conditions, *fas1-8 wee1-1 sog1-1* plants were able to restore the root growth phenotype of *fas1-8* plants back to WT length, demonstrating that SOG1 totally accounts for the observed phenotypes (Fig. 6B). Moreover, in accordance with SOG1 being responsible for the secondary checkpoint activated in *fas1-8 wee1-1*, the *fas1-8 wee1-1 sog1-1* triple mutant showed the same HU hypersensitivity phenotype as *wee1-1* (Fig. 6B).

To corroborate these data on the cell cycle level, we measured cellular parameters within the epidermis of mature (21 days after sowing) leaves of *fas1-8 wee1-1* versus *fas1-8 wee1-1 sog1-1* plants. The small leaf phenotype of the *fas1 wee1* mutants was found to correlate with a reduction in epidermal cell number rather than cell size, again indicating cell cycle checkpoint activation (Fig 6C-E). This decrease in cell number can be attributed to SOG1-function, as in the *fas1 wee1 sog1* triple mutant epidermal cell number and leaf were restored to near WT levels (Fig. 6C and D).

### Influence of the *ATM/ATR* Checkpoint on the Stability of DNA Repeats in Telomeres and 45S rDNA

In *fas1* and *fas2* single mutants, the stability of a subset of repetitive DNA is affected, and telomeres and rDNA are progressively lost (Mozgová et al., 2010). In agreement with the absence of any phenotypic differences between *fas1* and *fas1 wee1*, both genotypes displayed an equal reduction in rDNA copy number and telomere length (Fig. 7A and Fig. 8A).

**Figure 7.**
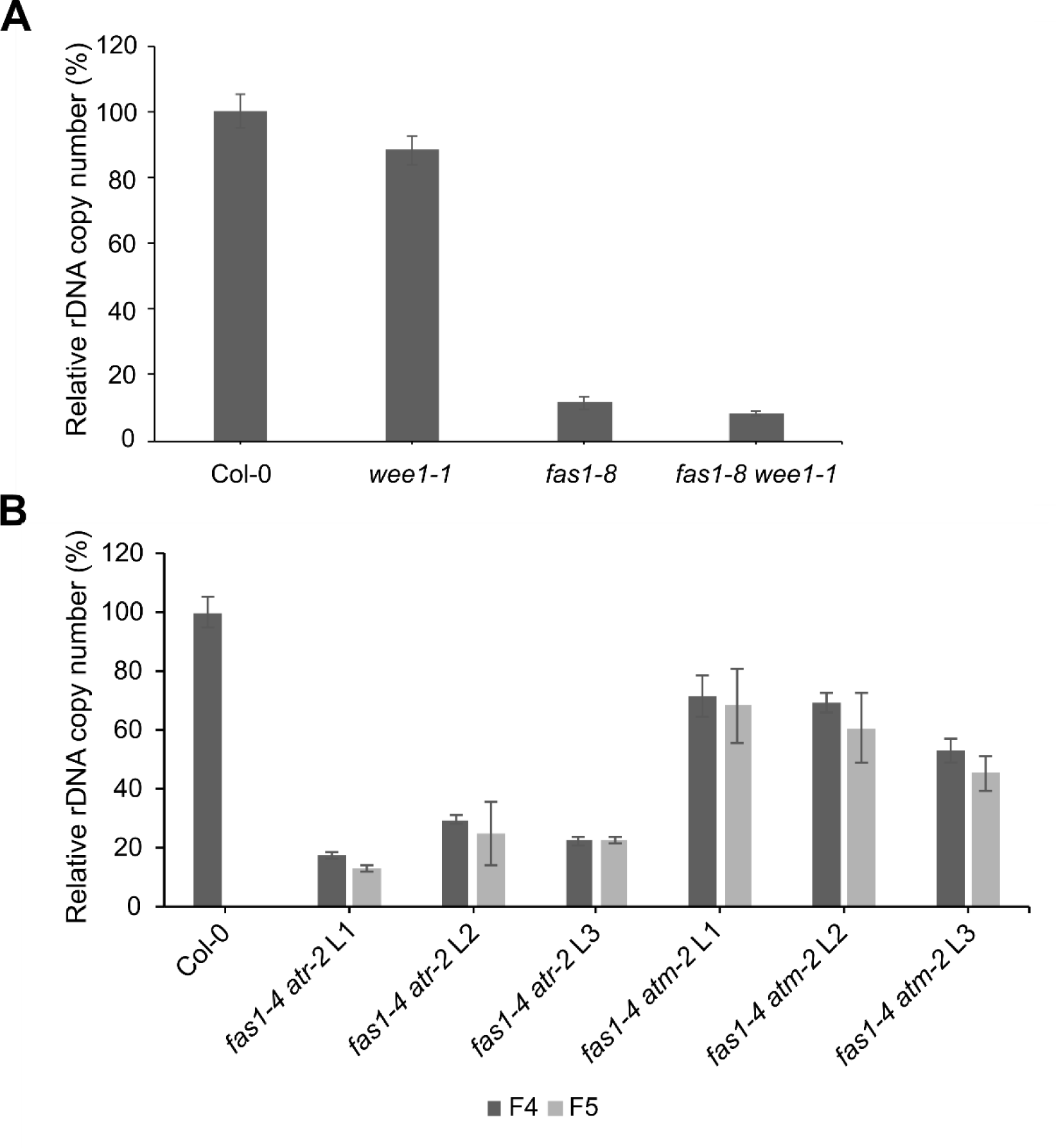
The loss of rDNA is suppressed in *fas1-4 atm-2*. qPCR analysis of relative rDNA copy number in WT and mutant lines. The 2^-ΔΔCT^ method was used, with 18S primers and *UBQ10* as reference. (A) rDNA copy numbers in WT (Col-0), *wee1-1, fas1-8* and *fas1-8 wee1-1*. Error bars in WT represent the SD between technical replicates, while in mutant lines, error bars correspond to the SD between biological replicates (n = 3). (B) rDNA copy numbers in *fas1 atm* and *fas1 atr* lines. F4 and F5 represent two consecutive generations of individual mutant lines, (L1-3). Error bars in WT and F4 generation represent the SD between technical replicates, since only one plant was used in F4 as a mother for several F5 plants. Error bars in F5 correspond to the SD between biological replicates, representing a progeny of F4 plant (n = 3).

**Figure 8.**
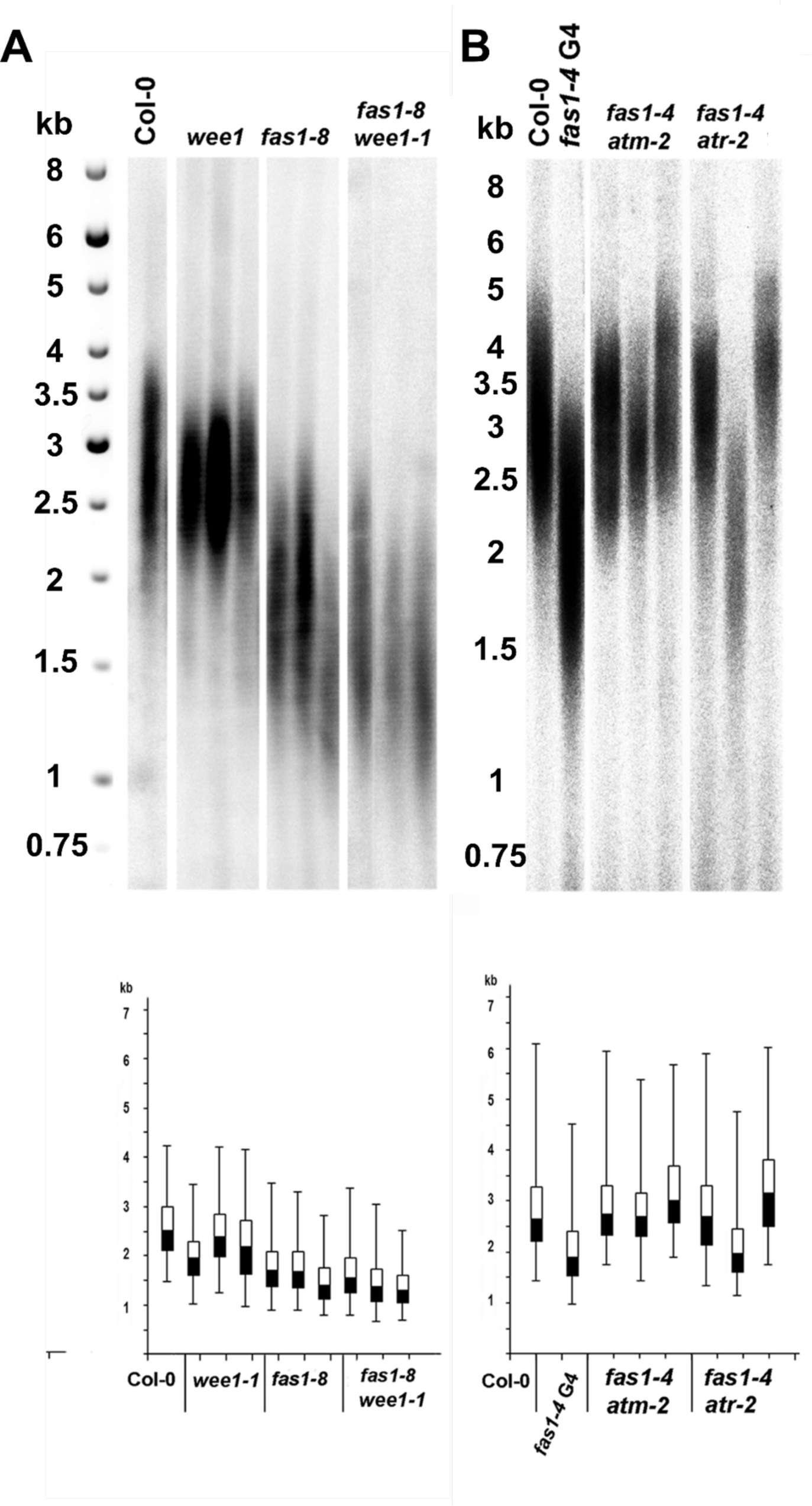
Mutation in *ATM* rescues the telomere shortening in *fas1*. Terminal restriction fragment analysis (TRF) in (A) Col-0, *wee1-1*, *fas1-8* and *fas1-8 wee1-1* double mutant and (B) *fas1-4 atm-2* and *fas1-4 atr-2* double mutants. Molecular markers (kb) are shown on the left, signal quantification of telomere lengths is shown in the bottom part of the pictures.

Subsequently, as both above and previously published data (Hisanaga et al., 2013) hint to a strong activation of ATM in the *fas1* mutant background, it was tested whether the rDNA loss and telomere erosion phenotypes were linked with the activated ATM and ATR checkpoints. In the *fas1 atm* plants, the systemic loss of rDNA was suppressed, but not to WT levels (Fig. 7B). Since these plants were obtained and analyzed in 4^th^ and 5^th^ filial generations, the results represent the steady-state levels of rDNA and not the immediate effect of loss of ATM function. Therefore, we investigated the rDNA change in the segregating offspring of a heterozygous *FAS1/fas1 ATM/atm* plant (Supplemental Fig. S4). The data showed that the *fas1 atm* double mutant line undergoes an initial rDNA reduction in F2, which is thus ATM-independent (Supplemental Fig. S4), but no further loss of rDNA copies is seen when these plants are propagated to the next generations (Fig. 7A), in contrast to *fas1* plants that lose increasing amounts of rDNA (Mozgová et al., 2010). We thus can conclude that ATM activation seems to be the predominant cause of the rDNA reduction in *fas1* mutants. Indeed, analysis of total rDNA copy numbers revealed a very low number in *fas1-4 atr-2* double mutants, comparable to the late generations of *fas1* mutants (Fig. 7B), confirming that the ATR pathway does not contribute to the rDNA phenotype in *fas1*.

We next analyzed the telomere length in *fas1 atm* and *fas1 atr* mutant lines (Fig. 8B, Supplemental Fig. S5). It has been shown previously that ATM and ATR regulate telomere length in a distinct manner, ATR by facilitating telomere replication, whereas ATM protects the chromosomal ends (Vespa et al., 2005; Amiard et al., 2011). Different results were observed in the individual double mutant lines. The *fas1 atm* line showed recovery of telomeres to WT levels (Fig. 8B), while in *fas1 atr* line-dependent heterogeneity emerged (Fig. 8B, Supplemental Fig. S5). From these results, we suggest that downstream targets of ATM mediate accelerated telomere shortening. The role of the ATR kinase is more complex and seems to be separated from WEE1 in terms of telomere processing.

## DISCUSSION

### Absence of CAF-1 Function Results in a G2/M Arrest, Making the S-Phase Checkpoint Redundant

Whereas in yeast and mammalian systems the *WEE1* kinase is essential for normal cell cycle progression, in plants its role is restricted to safeguarding the DNA replication phase (De Schutter et al., 2007; Cools et al., 2011). Here, we have identified a suppressor mutation in the *FAS1* gene that can partially rescue the hypersensitivity of *WEE1*^KO^ plants to replication stress. In contrast to the previously reported *trd1* suppressor mutation that rescues both cell death and decrease in meristem size of the *wee1* mutant upon replication stress (Kalhorzadeh et al., 2014), the *fas1* mutation rather appears to be epistatic to *wee1*. Although the roots of *fas1* mutants are already smaller compared to control plants, the *fas1 wee1* double mutant does not display the HU hypersensitivity phenotype of the *wee1* single mutant, indicating that the function of WEE1 is not required to cope with replication defects in *fas1* mutant roots. We demonstrate that this is likely caused by the activation of a G2/M checkpoint, resulting in a longer cell cycle. Contrary to previously reported (Schönrock et al., 2006), we found no difference in S-phase duration between control and *fas1* mutant plants. The reason for this discrepancy might be the different methods used: while in this work we measured the kinetics of DNA replication, the earlier report was based on a computational analysis of a single time point of a transcriptome study. Thus, even though CAF-1 function is mainly restricted to the DNA replication phase where it places histone H3-H4 dimers on the DNA strands, its absence apparently does not cause a delay in the S-phase. Nevertheless, an increase in S-phase duration is observed in *fas1-8 wee1-1* versus *fas1-8* plants, suggesting that WEE1 participates in the control of replication fork kinetics in the absence of FAS1, without however measurably interfering with cell cycle kinetics. This is different from what is observed for HU treatment, which resulted in a WEE1-dependent increase in S-phase duration in both WT and *fas1-8* mutant plants. This indicates that compared to impaired nucleosome assembly, nucleotide depletion exerts a much stronger effect on the replication process.

Histone loading in the *fas1* mutant is performed by the replication-independent pathway, which in Arabidopsis involves the HIRA complex (Duc et al., 2015), leading to the lower chromatin compactness and increased level of H3.3 (Kolarova et al., 2020). Because the genome will exist in a less compact state until the histones are loaded, it will thus accumulate more DSBs, which is evidenced by increased γH2AX foci in the *fas1* mutant (Muchová et al., 2015; Varas et al., 2017). Another possible explanation for the increased number of DSBs is that the nucleosomes containing histone H3.3 instead of histone H3.1, a variant that is not amenable to heterochromatic posttranslational modifications and thus cannot undergo compaction into heterochromatin (Jacob et al., 2014).

Double mutant analysis demonstrated that the prolonged cell cycle duration is predominantly due to activation of ATM activity, because the total cell cycle length was decreased in *fas1-4 atm-2* versus *fas1-4* mutants, despite an increase in S-phase duration. The latter is likely due to the activation of ATR, because no such increase was seen in *fas1-4 atr-2*. The observed increase in total cell cycle duration in *fas1-4 atr-2* might therefore be due to the accumulation of replication defects that eventually result in the activation of an ATM-dependent cell cycle arrest. In conclusion, although it appears that both ATR and ATM are active in FAS1-deficient plants, ATM likely accounts for the G2/M checkpoint that makes the S-phase checkpoint regulators become non-essential.

While ATR and ATM are known to share downstream components, and the damage recognized by one can be converted in order to be recognized by the other, it is for the first time shown here in plants that the ATR- and WEE1-regulated S-phase checkpoint can be partially rescued by constitutive activation of the ATM- and SOG1-mediated G2/M-phase checkpoint. Although it may at first seem surprising that a defect in chromosome assembly can correct for a replication problem, the situation reflects well the situation in budding yeast, where a downstream checkpoint can substitute an aberrant earlier cell cycle checkpoint, such as e.g. rescue of the G2 checkpoint mutant *rad9* through application of a microtubule depolymerizing drug that probably induces a mitotic spindle checkpoint (Weinert and Hartwell, 1988). It highlights the robustness of the cell cycle, where the failure of one checkpoint is rescued by a downstream checkpoint, a putative safeguard mechanism for maintaining genome stability. From this viewpoint it would be interesting to compare the mutation rate of the *fas1-8 wee1-1* double mutant with that of the *fas1-8 wee1-1 sog1-1* triple mutant that appears to overcome the *fas1-8*-induced growth arrest completely because of lack of both an intra-S phase and G2 checkpoint.

It is generally believed that the sensitivity of both *ATR*^KO^ and *WEE1*^KO^ plants to HU, as in other eukaryotic systems, mostly stems from the inability to coordinate replication firing with the reduced availability of dNTPs, resulting in replication fork stalling (Beck et al., 2012). In order to slow down replication firing, a decrease in S-phase CDK activity is necessary, which in Arabidopsis is mediated by WEE1 (De Schutter et al., 2007; Cools et al., 2011). It has been reported before that CDKA activity is decreased in the *fas1* mutant compared to the WT (Ramirez-Parra and Gutierrez, 2007). Given the lack of a phenotype of the *fas1 wee1* double mutant compared to the single *fas1* mutant, it seems unlikely that the WEE1 kinase largely contributes to this observed drop in CDK activity.

### ATM and ATR Activity Contributes to the Phenotypes of *fas1*

Whereas the impaired nucleosome assembly results in the activation of ATM and ATR, both indirectly contribute to the phenotype of the *fas1* mutant plants. Previously, we demonstrated that *fas1* mutants show a progressive telomere shortening and loss of 45S rDNA as plants go through mitosis (Mozgová et al., 2010). Here, we demonstrate that the lesions formed in rDNA in the *fas1* mutant are predominantly processed by ATM. This is in agreement with reports in mammals, which point to a unique role for ATM in the processing of DSBs in rDNA (Korsholm et al., 2019). Our previous data indicate that the process of rDNA recovery is rather stochastic, with large inter-individual variability between independently segregated plant lines (Pavlištová et al., 2016). Thus, knocking out ATM might be stopping recombinational processes causing dynamic changes in rDNA copy number (Pavlištová et al., 2016).

Telomere maintenance by ATM and ATR, as revealed here in the *fas1* background, is not as clear as their roles in rDNA stability. Telomeres are particularly sensitive to replication stress due to the end replication problem (Olovnikov, 1971), and compromised orchestration of factors involved in replication, or any imbalance during telomere replication leads to their instability, further detected as telomere shortening or elongation. Besides, during the DNA replication phase, cells with uncapped telomeres activate a DNA damage response, leading to chromosome fusions or formation of telomere dysfunction induced foci (Takai et al., 2003; Amiard et al., 2014). Thus, both ATR and ATM are known to participate in telomere maintenance and we show here that the *fas1* mutation is not an exception (Vespa et al., 2005; Amiard et al., 2011). While ATR facilitates replication of telomeres in cooperation with the CTC1-STN1-TEN1 (CST) complex, ATM recognizes the shortest telomeres and initiates recombination processes (Vespa et al., 2005; Amiard et al., 2011; Boltz et al., 2012). The exact mechanism of telomere erosion in *fas1* remains elusive. We recently demonstrated that telomere destabilization in *fas1* is not caused by a compromised telomerase function (Jaške et al., 2013), whose dysfunction would activate the alternative lengthening of telomeres, mediated through homologous recombination (Růčková et al., 2008). Disruption of ATM in *fas1* affects both types of repeats – rDNA as well as telomeres – since the activity of the DDR proteins is similarly deleterious for telomeres as it is for rDNA. Consistently, in *fas1 atr*, where telomeres show large inter-individual heterogeneity, ATM is still active and possibly responsible for this variability, independently of telomerase action.

### SOG1 Is Involved in the Increased Stress Tolerance Checkpoint

The question remains which kind of damage is perceived in the *fas1* mutants to activate the ATM-dependent DNA damage response. Is it the DSBs, that mostly affect rDNA and internal parts of the telomeres (Mozgová et al., 2010), which represent major repeated regions in the Arabidopsis genome and which are potential fragile sites (Dvořáčková et al., 2015); or is it the shortened telomeres, possibly falsely recognized as DSBs and also activating the DDR (Amiard et al., 2011)? Given that both *ATR*^KO^ and *WEE1*^KO^ in combination with *fas1* do not result in an amelioration of the *fas1* phenotype, it is tempting to speculate that the replication checkpoint is either not involved in activating the DDR or is only playing a minor role. In contrast, both ATM and SOG1 are clearly involved in the checkpoint activated in *fas1*, given that knocking them out confers a WT phenotype to *fas1* plants under control conditions. During replication stress, SOG1 plays a major role in the *fas1 wee1* mutant, as evidenced by the hypersensitive phenotype of *fas1 wee1 sog1* plants grown on HU. Although ATM is upstream of SOG1, the *fas1 wee1 atm* mutant is still resistant to HU, hinting at a possible role for ATR in activating SOG1 in these conditions. Our data lead to a possible model where mutation of *fas1* leads to telomere dysfunction and an increased number of DSBs, in this way activating the ATM pathway (Fig. 9). ATM activation leads to phosphorylation of downstream targets, including the SOG1 transcription factor. SOG1 will subsequently activate expression of (i) cell cycle regulators like *SMR7*, *ANAC044* and *ANAC085*, in this way possibly forcing a G2 arrest, premature exit of the mitotic cell cycle and/or entry into endoreplication, and (ii) DNA repair genes like RAD51 and its paralog RAD51B, which seem at least partly responsible for loss of rDNA copies (Muchová et al., 2015). The WT phenotype of *fas1* plants with a mutated *SOG1* gene shows that the induced G2-arrest is the cause of the macroscopic *fas1* phenotypes. Because of the activated ATM pathway in *fas1* mutants, it is very likely that some ATR targets are also activated, since the phosphorylation motif is identical, possibly explaining the higher resistance of *fas1 atr* plants to replication stress. Next to this, the G2/M checkpoint activated by ATM and SOG1 in *fas1 wee1* and *fas1 atr* plants likely allows the cell time to repair the damage caused by problems during the replication phase in these plants deficient for the intra-S-phase checkpoint (Fig. 9).

**Figure 9.**
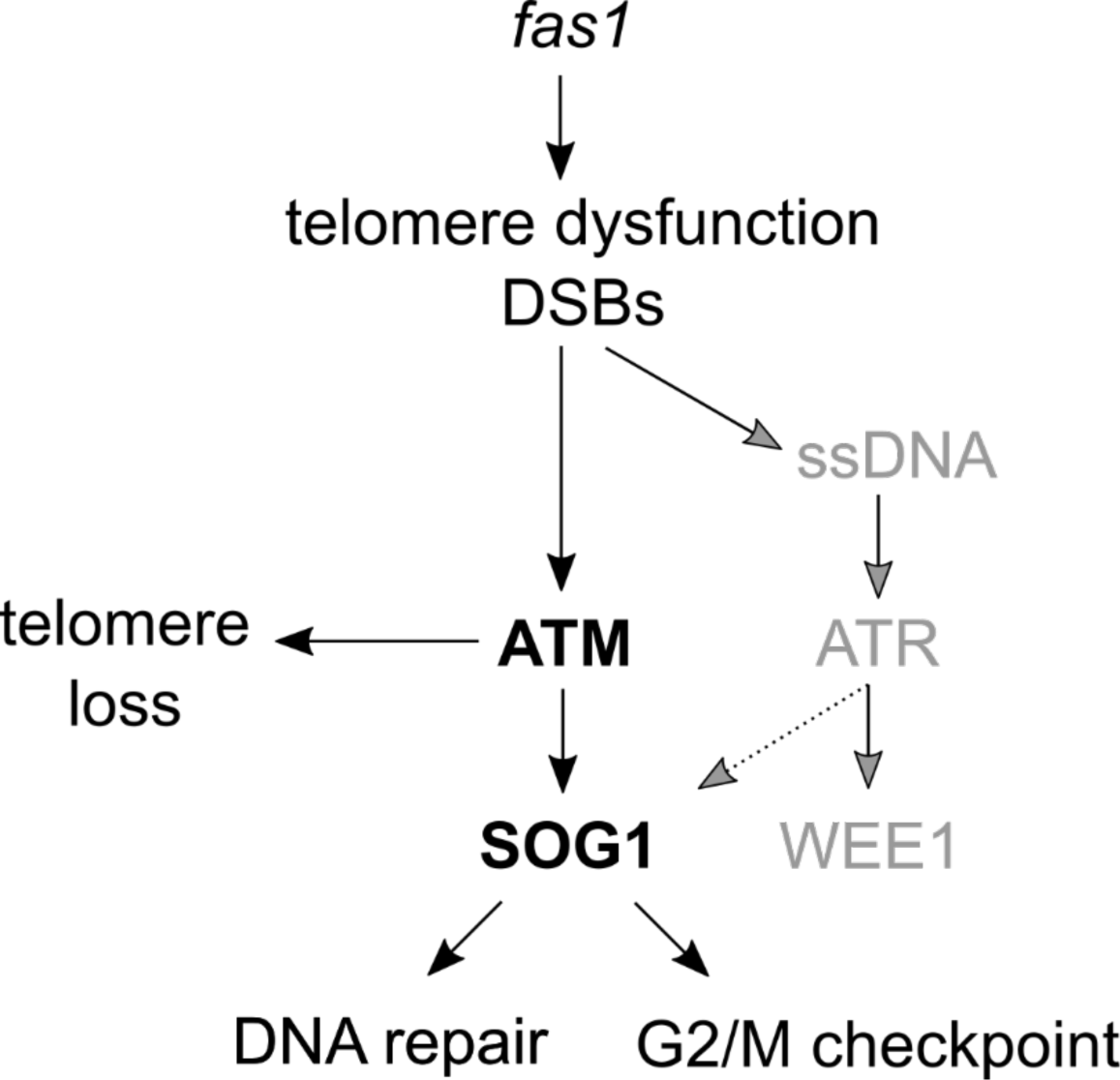
Possible model of activated DDR in *fas1* plants. The *FAS1* mutation leads to telomere dysfunction and formation of DSBs, in this way activating the ATM- and SOG1-dependent G2/M checkpoint. Activation of these DDR regulators overrules the need for intra-S-phase regulators (depicted in gray). For more details, see text.

## MATERIAL AND METHODS

### Plant Materials and Growth Conditions

*Arabidopsis thaliana* plants (all in Col-0 accession) were grown at 22°C under long-day conditions (16-h light/8 h dark) on half-strength Murashige & Skoog (MS) medium, 10 g/L sucrose, 0.5 g/L MES, pH 5.7 and 10 g/L agar (ref). The *fas1-8* (GK_095A01) allele was acquired from GABI-Kat. The *atm-2*, *atr-2*, *wee1-1, fas1-4*, and *sog1-1* alleles were described previously (Garcia et al., 2003; Culligan et al., 2004; Exner et al., 2006; De Schutter et al., 2007; Yoshiyama et al., 2009; Hisanaga et al., 2013). Homozygous insertion alleles were confirmed by genotyping PCR using primers listed in Supplemental Table S1. For treatment with HU, seeds were directly plated on control medium and medium containing 0.25, 0.5 or 0.75 mM HU. The root length of 7-d-old plants was measured. For treatment with zeocin, seeds were directly plated on control medium, medium containing 12 µM zeocin and medium containing 12 µM zeocin and 0.5 mM HU. For treatment with bleomycin, seeds were sown on sterilized membranes on ½ MS medium. After 2d of germination and 4d of growth, the membrane was transferred to ½ MS medium and ½ MS medium containing 0.6 µg/ml bleomycin for 24h.

### Meta-Analysis of Transcriptomes

Three studies were used for this meta-analysis: in the study of Schönrock, ATH1 microarrays were used to study differential gene expression of whole *fas1* and *fas2* seedlings at 12 DAS (Schönrock et al., 2006). In the Hisanaga study, ATH1 microarrays were used on the leaf primordia of *fas1-5* plants at 10 DAS (Hisanaga et al., 2013), whereas the Mozgová study used AGRONOMICS1 tiling arrays on soil-grown *fas2-4* plants at 48 DAS (Mozgová et al., 2015). From these three different studies, the genes upregulated with an FDR < 0.05 and FC > 1.5 were compared. The overlap was then compared to a list of 61 genes classified as DDR markers (Yi et al., 2014).

### EMS Mutagenesis and Mapping

EMS mutagenesis was performed as described in Kalhorzadeh et al. (2014). Briefly, *wee1-1* seeds were treated with EMS and sown in 200 pools of 250 seeds each. After self-fertilization, M1 seeds were grown and M2 seeds were collected from individual M1 plants. M2 plants were then screened for increased root growth on HU compared to *wee1-1* plants. Line 49-2 showed increased resistance to HU and was crossed to the Landsberg *erecta* accession containing the *wee1-1* allele. F1 plants were allowed to self-fertilize and F2 plants were again screened for increased resistance to HU. Leaf samples of about 200 plants were pooled and nuclear DNA was extracted with a DNeasy Plant Mini Kit (QIAGEN) according to the manufacturer’s protocol. Illumina Tru-Seq libraries were generated according to the manufacturer’s protocol and sequenced on an Illumina NextSeq 500 150-bp paired-end run. To align the reads to the reference genome (Col-0; TAIR10), the SHORE pipeline was used (Ossowski et al., 2008). Based on the alignment, single-nucleotide polymorphisms were determined and relative allele frequencies were compared between the two parental genomes (Col-0 and L*er*) using SHOREmap (Schneeberger et al., 2009).

### Flow Cytometry, Microscopy and Confocal Microscopy

To obtain ploidy profiles, leaf material of 21-d-old plants was chopped in 200 µL Cystain UV Precise P nuclei extraction buffer (Partec), supplemented with 800 µL staining buffer, filtered and measured by a Cyflow MB flow cytometer (Partec).

For leaf measurements, first leaves were harvested at 21 d after sowing on control medium, cleared overnight in ethanol, stored in lactic acid for microscopy, and observed with a microscope fitted with differential interference contrast optics (Leica DMLB). The total (blade) area was determined from images digitized directly with a digital camera mounted on a stereozoom microscope (Stemi SV11; Zeiss). From scanned drawing-tube images of the outlines of at least 30 cells of the abaxial epidermis located between 25 to 75% of the distance between the tip and the base of the leaf, halfway between the midrib and the leaf margin, the following parameters were determined: total area of all cells in the drawing and total numbers of pavement and guard cells, from which the average cell area was calculated. The total number of cells per leaf was estimated by dividing the leaf area by the average cell area (De Veylder et al., 2001). Leaf size, epidermal cell number and epidermal cell size in the different lines were analyzed and compared by mixed model analysis (SAS Studio).

To visualize the root meristems, roots were stained for 3 min in a 10-µM propidium iodide solution (Sigma-Aldrich) and were analyzed with an LSM 5 exciter confocal microscope (Zeiss).

### EdU Time Course

For cell cycle length analysis, we used a method adapted from Hayashi et al. (2013). Plants were grown on vitamin-supplemented MS medium (10 g/L sucrose, 0.1 g/L myo-inositol, 0.5 g/L MES, 100 μL thiamine hydrochloride (10 mg/mL), 100 μL pyridoxine (5 mg/mL), 100 μL nicotinic acid (5 mg/mL), pH 5.7, adjusted with 1 m KOH, and 10 g/L agar) for 5 days, and transferred to the same medium supplemented with EdU (10 µM). Samples were collected after 3, 6, 9 and 12 h, fixed in paraformaldehyde (4% in PME buffer: 50 mm piperazine-N,N′-bis(2-ethanesulphonic acid) (PIPES), pH 6.9; 5 mM MgSO_4_; 1 mM EGTA) for 45 min and washed with PME 1X buffer. Root apices were dissected on a glass slide and digested in a drop of enzyme mix (1% (w/v) cellulase, 0.5% (w/v) cytohelicase, 1% (w/v) pectolyase in PME) for 1 h at 37°C. After three washes with PME 1X, root apices were squashed gently between the slide and a coverslip, and frozen in liquid nitrogen. After removal of the coverslip and drying of the slides overnight, EdU revelation and Hoechst counterstaining were performed following the Click-iT Plus EdU Alexa Fluor 488 Imaging Kit instructions (Thermo Fisher Scientific). The percentage of EdU-positive nuclei was plotted as a function of time. The percentage of EdU-positive nuclei increases linearly with time, and follows an equation that can be written as y = at + b, in which y is the percentage of EdU-positive nuclei and t is time. Total cell cycle length is estimated as 100/a, and S-phase length is b/a.

### EdU Pulse Labeling

Progression through G2/M of WT and mutant lines was analyzed using pulse labeling with EdU. Briefly, 5-d-old seedlings were grown on MS plates, then transferred to liquid MS medium with 10-µM EdU, followed by a 15-min incubation. After washing with MS medium, seedlings were transferred again to MS medium and collected after 4, 8, and 12 h. Later, nuclei were stained with DAPI and cells with mitotic figures were counted. Data are presented as the percentage of EdU-labelled cells among those with mitotic figures.

### Terminal Restriction Fragment (TRF) Analyses

TRF analysis was performed as described in Mozgová et al. (2010). Plant DNA was extracted from 5-w-old leaves in accordance with Dellaporta et al. (1983) and 400 ng of gDNA was used for the analysis. DNA was digested with 20 U of MseI (NEB) at 37°C overnight, DNA fragments were separated on the 0.8% agarose gel and alkali-blotted onto a Hybond XL membrane (GE Healthcare Life Sciences, Waukesha, WI, USA). Telomere fragments were detected by a radioactively labeled telomeric DNA probe, prepared by non-template PCR according to Ijdo et al. (1991) and labelled with [α-^32^P] ATP using random decamers (Decalabel DNA labelling kit, Thermo Fisher Scientific). Hybridization was performed overnight at 65°C. Telomeric signals were visualized using a FLA7000 imager (Fujifilm) and quantified as described in Zachová et al. (2013). Quantification was carried out as follows: the medians of TRF lengths were calculated as Σ(ODi x Li)/Σ(ODi), in which ODi is the signal intensity above the background within intervals i, and Li is the molecular weight (kb) at the midpoint of interval I. In the charts, the bottom and top of the box represent the first and third quartile, respectively, separated by the median, and s.d. intervals above and under the box are the maximum and minimum of the telomere lengths, respectively.

### Determination of rDNA Copy Number by qPCR

Quantitative PCR (qPCR) analysis of 45S rDNA copy number qPCR was performed in three technical and three biological repeats, using primers for 18S region and normalized to *UBIQUITIN 10* under the following conditions: initial denaturation 95°C/7 min, 35 cycles of 95°C/30 s, 56°C/30 s, 72°C/30 s with final incubation at 75°C/5 min followed by the standard melting analysis. The analysis was performed by StepOnePlusTM Real-Time PCR system (Applied Biosystems) using FastStart SYBR GreenMaster (Roche, http://www.roche.com/).

### qRT-PCR Analysis

RNA was extracted from root tips (1-2 mm) of 7-d-old seedlings using the RNEasy plant mini kit (QIAGEN), according to the manufacturer’s protocol. cDNA was prepared from 1 µg of RNA using iScript cDNA synthesis kit (Bio-Rad). Quantitative RT-PCR was performed with Lightcycler 480 SYBR Green I Master (Roche) in a total volume of 5 µL and analyzed on a Lightcycler 480 (Roche). Each reaction was done in three technical and three biological repeats. Expression levels of each gene were normalized to the following reference genes: *EMB2386*, *RPS26C* and *PAC1*. Primers used for qRT-PCR are given in Supplemental Table S1.

### Accession Numbers

FAS1: AT1G65470; ATM: AT3G48190; ATR: AT5G40820; SOG1: AT1G25580; WEE1: AT1G02970; SMR7: AT3G27630

## Supplemental Data

The following supplemental materials are available.

**Supplemental Figure S1. Confirmation of the mutation in *FAS1* in line 49-2 by Sanger sequencing**

**Supplemental Figure S2. 49-2 revertant phenocopies the *fas1-8* knockout line.**

**Supplemental Figure S3. Treatment with a DNA damaging agent induces DDR genes to a similar extent in *fas1* background as in WT.**

**Supplemental Figure S4. rDNA copy number of plants segregated from heterozygous *FAS1/fas1 ATM/atm* plants.**

**Supplemental Figure S5. Terminal restriction fragment analysis of additional lines.**

**Supplemental Table S1. Cell cycle parameters of Col-0, *wee1-1, fas1-8* and *fas1-8 wee1-1* under control conditions.**

**Supplemental Table S2. Cell cycle parameters of Col-0, *wee1-1, fas1-8* and *fas1-8 wee1-1* under replication stress.**

**Supplemental Table S3. Cell cycle parameters of Col-0, *fas1-4, fas1-4 atm-2, fas1-4 atr-2, atr-2* and *atm-2* under control conditions.**

**Supplemental Table S4. List of primers used for genotyping and qRT-PCR.**

## ACKNOWLEDGMENTS

The authors thank Annick Bleys for help in preparing the manuscript and Dr. Tetsuya Hisanaga (GMI, Vienna) for *fas1 atr* and *fas1 atm* mutants. Core Facility Plants Sciences of CEITEC MU is gratefully acknowledged for the obtaining of the scientific data presented in this paper.

## Reference

Amiard, S., Da Ines, O., Gallego, M.E., and White, C.I. (2014). Responses to telomere erosion in plants. PLoS ONE 9, e86220.

Amiard, S., Depeiges, A., Allain, E., White, C.I., and Gallego, M.E. (2011). *Arabidopsis* ATM and ATR kinases prevent propagation of genome damage caused by telomere dysfunction. Plant Cell 23, 4254–4265.

Beck, H., Nähse-Kumpf, V., Larsen, M.S.Y., O’Hanlon, K.A., Patzke, S., Holmberg, C., Mejlvang, J., Groth, A., Nielsen, O., Syljuåsen, R.G., and Sørensen, C.S. (2012). Cyclin-dependent kinase suppression by WEE1 kinase protects the genome through control of replication initiation and nucleotide consumption. Mol. Cell. Biol. 32, 4226–4236.

Boltz, K.A., Leehy, K., Song, X., Nelson, A.D., and Shippen, D.E. (2012). ATR cooperates with CTC1 and STN1 to maintain telomeres and genome integrity in *Arabidopsis*. Mol. Biol. Cell 23, 1558–1568.

Bourbousse, C., Vegesna, N., and Law, J.A. (2018). SOG1 activator and MYB3R repressors regulate a complex DNA damage network in *Arabidopsis*. Proc. Natl. Acad. Sci. USA 115, E12453–E12462.

Cools, T., Iantcheva, A., Weimer, A.K., Boens, S., Takahashi, N., Maes, S., Van den Daele, H., Van Isterdael, G., Schnittger, A., and De Veylder, L. (2011). The *Arabidopsis thaliana* checkpoint kinase WEE1 protects against premature vascular differentiation during replication stress. Plant Cell 23, 1435–1448.

Culligan, K., Tissier, A., and Britt, A. (2004). ATR regulates a G2-phase cell-cycle checkpoint in *Arabidopsis thaliana*. Plant Cell 16, 1091–1104.

De Schutter, K., Joubès, J., Cools, T., Verkest, A., Corellou, F., Babiychuk, E., Van Der Schueren, E., Beeckman, T., Kushnir, S., Inzé, D., and De Veylder, L. (2007). *Arabidopsis* WEE1 kinase controls cell cycle arrest in response to activation of the DNA integrity checkpoint. Plant Cell 19, 211–225.

De Veylder, L., Beeckman, T., Beemster, G.T., Krols, L., Terras, F., Landrieu, I., van der Schueren, E., Maes, S., Naudts, M., and Inze, D. (2001). Functional analysis of cyclin-dependent kinase inhibitors of Arabidopsis. Plant Cell 13, 1653–1668.

Dellaporta, S.L., Wood, J., and Hicks, J.B. (1983). A plant DNA minipreparation: version II. Plant Mol. Biol. Rep. 1, 19–21.

Duc, C., Benoit, M., Le Goff, S., Simon, L., Poulet, A., Cotterell, S., Tatout, C., and Probst, A.V. (2015). The histone chaperone complex HIR maintains nucleosome occupancy and counterbalances impaired histone deposition in CAF-1 complex mutants. Plant J. 81, 707–722.

Dvořáčková, M., Fojtová, M., and Fajkus, J. (2015). Chromatin dynamics of plant telomeres and ribosomal genes. Plant J. 83, 18–37.

Eekhout, T., Kalhorzadeh, P., and De Veylder, L. (2015). Lack of RNase H2 activity rescues HU-sensitivity of WEE1 deficient plants. Plant Signal. Behav. 10, e1001226.

Endo, M., Ishikawa, Y., Osakabe, K., Nakayama, S., Kaya, H., Araki, T., Shibahara, K.-i., Abe, K., Ichikawa, H., Valentine, L., Hohn, B., and Toki, S. (2006). Increased frequency of homologous recombination and T-DNA integration in *Arabidopsis* CAF-1 mutants. EMBO J. 25, 5579–5590.

Exner, V., Taranto, P., Schönrock, N., Gruissem, W., and Hennig, L. (2006). Chromatin assembly factor CAF-1 is required for cellular differentiation during plant development. Development 133, 4163–4172.

Garcia, V., Bruchet, H., Camescasse, D., Granier, F., Bouchez, D., and Tissier, A. (2003). *AtATM* is essential for meiosis and the somatic response to DNA damage in plants. Plant Cell 15, 119–132.

Hayashi, K., Hasegawa, J., and Matsunaga, S. (2013). The boundary of the meristematic and elongation zones in roots: endoreduplication precedes rapid cell expansion. Sci. Rep. 3, 2723.

Hisanaga, T., Ferjani, A., Horiguchi, G., Ishikawa, N., Fujikura, U., Kubo, M., Demura, T., Fukuda, H., Ishida, T., Sugimoto, K., and Tsukaya, H. (2013). The ATM-dependent DNA damage response acts as an upstream trigger for compensation in the *fas1* mutation during Arabidopsis leaf development. Plant Physiol. 162, 831–841.

Ijdo, J.W., Wells, R.A., Baldini, A., and Reeders, S.T. (1991). Improved telomere detection using a telomere repeat probe (TTAGGG) n generated by PCR. Nucleic Acids Res. 19, 4780.

Jacob, Y., Bergamin, E., Donoghue, M.T.A., Mongeon, V., LeBlanc, C., Voigt, P., Underwood, C.J., Brunzelle, J.S., Michaels, S.D., Reinberg, D., Couture, J.-F., and Martienssen, R.A. (2014). Selective methylation of histone H3 variant H3.1 regulates heterochromatin replication. Sci. Adv. 343, 1249–1253.

Jaške, K., Mokroš, P., Mozgová, I., Fojtová, M., and Fajkus, J. (2013). A telomerase-independent component of telomere loss in chromatin assembly factor 1 mutants of *Arabidopsis thaliana*. Chromosoma 122, 285–293.

Kalhorzadeh, P., Hu, Z., Cools, T., Amiard, S., Willing, E.-M., De Winne, N., Gevaert, K., De Jaeger, G., Schneeberger, K., White, C.I., and De Veylder, L. (2014). *Arabidopsis thaliana* RNase H2 deficiency counteracts the needs for the WEE1 checkpoint kinase but triggers genome instability. Plant Cell 26, 3680–3692.

Kaufman, P.D., Kobayashi, R., and Stillman, B. (1997). Ultraviolet radiation sensitivity and reduction of telomeric silencing in *Saccharomyces cerevisiae* cells lacking chromatin assembly factor-I. Genes Dev. 11, 345–357.

Kaufman, P.D., Kobayashi, R., Kessler, N., and Stillman, B. (1995). The p150 and p60 subunits of chromatin assembly factor-I: a molecular link between newly synthesized histones and DNA replication. Cell 81, 1105–1114.

Kaya, H., Shibahara, K.-i., Taoka, K.-i., Iwabuchi, M., Stillman, B., and Araki, T. (2001). *FASCIATA* genes for chromatin assembly factor-1 in *Arabidopsis* maintain the cellular organization of apical meristems. Cell 104, 131–142.

Kirik, A., Pecinka, A., Wendeler, E., and Reiss, B. (2006). The chromatin assembly factor subunit FASCIATA1 is involved in homologous recombination in plants. Plant Cell 18, 2431–2442.

Kolarova, K., Dadejova, M.N., Loja, T., Lochmanova, G., Sykorova, E., and Dvorackova, M. (2020). Disruption of NAP1 genes in Arabidopsis thaliana suppresses the fas1 mutant phenotype, enhances genome stability and changes the chromatin compaction. Plant J.

Korsholm, L.M., Gál, Z., Lin, L., Quevedo, O., Ahmad, D.A., Dulina, E., Luo, Y., Bartek, J., and Larsen, D.H. (2019). Double-strand breaks in ribosomal RNA genes activate a distinct signaling and chromatin response to facilitate nucleolar restructuring and repair. Nucleic Acids Res. 47, 8019–8035.

Leyser, H.M.O., and Furner, I.J. (1992). Characterization of three shoot apical meristem mutants of *Arabidopsis thaliana*. Development 116, 397–403.

Mozgova, I., Mokros, P., and Fajkus, J. (2010). Dysfunction of chromatin assembly factor 1 induces shortening of telomeres and loss of 45S rDNA in Arabidopsis thaliana. Plant Cell 22, 2768–2780.

Mozgova, I., Wildhaber, T., Trejo-Arellano, M.S., Fajkus, J., Roszak, P., Kohler, C., and Hennig, L. (2018). Transgenerational phenotype aggravation in CAF-1 mutants reveals parent-of-origin specific epigenetic inheritance. New Phytol 220, 908–921.

Mozgová, I., Mokroš, P., and Fajkus, J. (2010). Dysfunction of chromatin assembly factor 1 induces shortening of telomeres and loss of 45S rDNA in *Arabidopsis thaliana*. Plant Cell 22, 2768–2780.

Mozgová, I., Wildhaber, T., Liu, Q., Abou-Mansour, W., L’Haridon, F., Metraux, J.-P., Gruissem, W., Hofius, D., and Hennig, L. (2015). Chromatin assembly factor CAF-1 represses priming of plant defence response genes. Nat. Plants 1, 15127.

Muchová, V., Amiard, S., Mozgová, I., Dvořáčková, M., Gallego, M.E., White, C., and Fajkus, J. (2015). Homology-dependent repair is involved in 45S rDNA loss in plant CAF-1 mutants. Plant J. 81, 198–209.

Ogita, N., Okushima, Y., Tokizawa, M., Yamamoto, Y.Y., Tanaka, M., Seki, M., Makita, Y., Matsui, M., Okamoto-Yoshiyama, K., Sakamoto, T., Kurata, T., Hiruma, K., Saijo, Y., Takahashi, N., and Umeda, M. (2018). Identifying the target genes of SUPPRESSOR OF GAMMA RESPONSE 1, a master transcription factor controlling DNA damage response in *Arabidopsis*. Plant J. 94, 439–453.

Olovnikov, A.M. (1971). Principle of marginotomy in template synthesis of polynucleotides. Dokl. Akad. nauk SSSR 201, 1496–1499.

Ossowski, S., Schneeberger, K., Clark, R.M., Lanz, C., Warthmann, N., and Weigel, D. (2008). Sequencing of natural strains of *Arabidopsis thaliana* with short reads. Genome Res. 18, 2024–2033.

Pan, T., Qin, Q., Nong, C., Gao, S., Wang, L., Cai, B., Zhang, M., Wu, C., Chen, H., Li, T., Xiong, D., Li, G., Wang, S., and Yan, S. (2021). A novel WEE1 pathway for replication stress responses. Nat Plants.

Pavlištová, V., Dvořáčková, M., Jež, M., Mozgová, I., Mokroš, P., and Fajkus, J. (2016). Phenotypic reversion in *fas* mutants of *Arabidopsis thaliana* by reintroduction of *FAS* genes: variable recovery of telomeres with major spatial rearrangements and transcriptional reprogramming of 45S rDNA genes. Plant J. 88, 411–424.

Ramirez-Parra, E., and Gutierrez, C. (2007). E2F regulates *FASCIATA1*, a chromatin assembly gene whose loss switches on the endocycle and activates gene expression by changing the epigenetic status. Plant Physiol. 144, 105–120.

Roldán-Arjona, T., and Ariza, R.R. (2009). Repair and tolerance of oxidative DNA damage in plants. Mutat. Res.-Rev. Mutat. Res. 681, 169–179.

Růčková, E., Friml, J., Procházková Schrumpfová, P., and Fajkus, J. (2008). Role of alternative telomere lengthening unmasked in telomerase knock-out mutant plants. Plant Mol. Biol. 66, 637–646.

Schneeberger, K., Ossowski, S., Lanz, C., Juul, T., Petersen, A.H., Nielsen, K.L., Jørgensen, J.-E., Weigel, D., and Andersen, S.U. (2009). SHOREmap: simultaneous mapping and mutation identification by deep sequencing. Nat. Methods 6, 550–551.

Schönrock, N., Exner, V., Probst, A., Gruissem, W., and Hennig, L. (2006). Functional genomic analysis of CAF-1 mutants in *Arabidopsis thaliana*. J. Biol. Chem. 281, 9560–9568.

Serra, H., Da Ines, O., Degroote, F., Gallego, M.E., and White, C.I. (2013). Roles of XRCC2, RAD51B and RAD51D in RAD51-independent SSA recombination. PLoS Genet. 9, e1003971.

Sjogren, C.A., Bolaris, S.C., and Larsen, P.B. (2015). Aluminum-dependent terminal differentiation of the Arabidopsis root tip is mediated through an ATR-, ALT2-, and SOG1-regulated transcriptional response. Plant Cell 27, 2501-2515.

Smith, S., and Stillman, B. (1989). Purification and characterization of CAF-I, a human cell factor required for chromatin assembly during DNA replication in vitro. Cell 58, 15–25.

Takahashi, N., Ogita, N., Takahashi, T., Taniguchi, S., Tanaka, M., Seki, M., and Umeda, M. (2019). A regulatory module controlling stress-induced cell cycle arrest in *Arabidopsis*. eLife 8, e43944.

Takai, H., Smogorzewska, A., and de Lange, T. (2003). DNA damage foci at dysfunctional telomeres. Curr. Biol. 13, 1549–1556.

Varas, J., Santos, J.L., and Pradillo, M. (2017). The absence of the Arabidopsis chaperone complex CAF-1 produces mitotic chromosome abnormalities and changes in the expression profiles of genes involved in DNA repair. Front. Plant Sci. 8, 525.

Verreault, A., Kaufman, P.D., Kobayashi, R., and Stillman, B. (1996). Nucleosome assembly by a complex of CAF-1 and acetylated histones H3/H4. Cell 87, 95–104.

Vespa, L., Couvillion, M., Spangler, E., and Shippen, D.E. (2005). ATM and ATR make distinct contributions to chromosome end protection and the maintenance of telomeric DNA in *Arabidopsis*. Genes Dev. 19, 2111–2115.

Wang, L., Zhan, L., Zhao, Y., Huang, Y., Wu, C., Pan, T., Qin, Q., Xu, Y., Deng, Z., Li, J., Hu, H., Xue, S., and Yan, S. (2021). The ATR-WEE1 kinase module inhibits the MAC complex to regulate replication stress response. Nucleic Acids Res.

Weinert, T.A., and Hartwell, L.H. (1988). The RAD9 gene controls the cell cycle response to DNA damage in Saccharomyces cerevisiae. Science 241, 317–322.

Yi, D., Lessa Alvim Kamei, C., Cools, T., Vanderauwera, S., Takahashi, N., Okushima, Y., Eekhout, T., Yoshiyama, K.O., Larkin, J., Van den Daele, H., Conklin, P., Britt, A., Umeda, M., and De Veylder, L. (2014). The *Arabidopsis* SIAMESE-RELATED cyclin-dependent kinase inhibitors SMR5 and SMR7 regulate the DNA damage checkpoint in response to reactive oxygen species. Plant Cell 26, 296–309.

Yoshiyama, K., Conklin, P.A., Huefner, N.D., and Britt, A.B. (2009). Suppressor of gamma response 1 (*SOG1*) encodes a putative transcription factor governing multiple responses to DNA damage. Proc. Natl. Acad. Sci. USA 106, 12843–12848.

Yoshiyama, K.O., Kobayashi, J., Ogita, N., Ueda, M., Kimura, S., Maki, H., and Umeda, M. (2013). ATM-mediated phosphorylation of SOG1 is essential for the DNA damage response in *Arabidopsis*. EMBO Rep. 14, 817–822.

Zachová, D., Fojtová, M., Dvořáčková, M., Mozgová, I., Lermontova, I., Peška, V., Schubert, I., Fajkus, J., and Sýkorová, E. (2013). Structure-function relationships during transgenic telomerase expression in *Arabidopsis*. Physiol. Plant. 149, 114–126.

Zhou, B.-B., and Elledge, S.J. (2000). The DNA damage response: putting checkpoints in perspective. Nature 408, 433–439.

Zhu, Y., Dong, A., Meyer, D., Pichon, O., Renou, J.-P., Cao, K., and Shen, W.-H. (2006). *Arabidopsis NRP1* and *NRP2* encode histone chaperones and are required for maintaining postembryonic root growth. Plant Cell 18, 2879–2892.

Zhu, Y., Weng, M., Yang, Y., Zhang, C., Li, Z., Shen, W.-H., and Dong, A. (2011). *Arabidopsis* homologues of the histone chaperone ASF1 are crucial for chromatin replication and cell proliferation in plant development. Plant J. 66, 443–455.

